# Individual differences in haemoglobin concentration influence BOLD fMRI functional connectivity and its correlation with cognition

**DOI:** 10.1101/835660

**Authors:** Phillip G. D. Ward, Edwina R. Orchard, Stuart Oldham, Aurina Arnatkevičiūtė, Francesco Sforazzini, Alex Fornito, Elsdon Storey, Gary F. Egan, Sharna D. Jamadar

## Abstract

Resting-state connectivity measures the temporal coherence of the spontaneous neural activity of spatially distinct regions, and is commonly measured using BOLD-fMRI. The BOLD response follows neuronal activity, when changes in the relative concentration of oxygenated and deoxygenated haemoglobin cause fluctuations in the MRI T2* signal. Since the BOLD signal detects changes in relative concentrations of oxy/deoxy-haemoglobin, individual differences in haemoglobin levels may influence the BOLD signal-to-noise ratio in a manner independent of the degree of neural activity. In this study, we examined whether group differences in haemoglobin may confound measures of functional connectivity. We investigated whether relationships between measures of functional connectivity and cognitive performance could be influenced by individual variability in haemoglobin. Finally, we mapped the neuroanatomical distribution of the influence of haemoglobin on functional connectivity to determine where group differences in functional connectivity are manifest.

In a cohort of 518 healthy elderly subjects (259 men) each sex group was median split into two groups with high and low haemoglobin concentration. Significant differences were obtained in functional connectivity between the high and low haemoglobin groups for both men and women (Cohen’s d 0.17 and 0.03 for men and women respectively). The haemoglobin connectome in males showed a widespread systematic increase in functional connectivity correlational scores, whilst the female connectome showed predominantly parietal and subcortical increases and temporo-parietal decreases. Despite the haemoglobin groups having no differences in cognitive measures, significant differences in the linear relationships between cognitive performance and functional connectivity were obtained for all 5 cognitive tests in males, and 4 out of 5 tests in females.

Our findings confirm that individual variability in haemoglobin levels that give rise to group differences are an important confounding variable in BOLD-fMRI-based studies of functional connectivity. Controlling for haemoglobin variability as a potentially confounding variable is crucial to ensure the reproducibility of human brain connectome studies, especially in studies that compare groups of individuals, compare sexes, or examine connectivity-cognition relationships.

**Highlights:**

- Individual differences in haemoglobin significantly impact measures of functional connectivity in the elderly.
- Significant differences in connectivity-cognition relationships are shown between groups separated by haemoglobin values without accompanying cognitive differences.
- The influence of haemoglobin on functional connectivity differs between men and women.

## 1. Introduction

Functional connectivity refers to statistical dependencies between spatially distinct neurophysiological signals (Friston et al., 1994), and can be estimated in humans at a whole-brain level using blood-oxygenation-level-dependent (BOLD) functional magnetic resonance imaging (fMRI). Many studies measure functional connectivity using task-free resting-state BOLD fMRI data (Biswal et al., 1995; Fox and Raichle, 2007), otherwise known as resting-state functional connectivity. Numerous studies have demonstrated that resting-state functional connectivity is associated with variability in cognition (Fox et al., 2007; Jamadar et al., 2016), age (Andrews-Hanna et al., 2007), sex (Jamadar et al., 2018; Weiss et al., 2003), genetics (Fornito et al., 2011; Glahn et al., 2010), psychiatric conditions (Fornito and Bullmore, 2010; Garrity et al., 2007; McGrath et al., 2013), and neurodegeneration (e.g., Alzheimer’s disease, (Greicius et al., 2004)).

Measures of functional connectivity depend on temporal fluctuations in the BOLD signal. Changes to the relative concentrations of oxygenated and deoxygenated haemoglobin in a brain region cause fluctuations to the MRI T2* signal, giving rise to the BOLD effect (Ogawa et al., 1992). The BOLD signal therefore provides an indirect measure of brain function arising from the neurovascular coupling between neuronal activity and cerebral haemodynamics (Phillips et al., 2016). As described by Liu (Liu, 2017), the relative signal change in BOLD-fMRI is given as

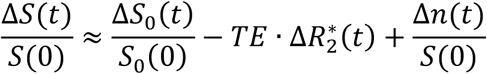

where *S(t)* denotes the signal acquired at time *t, R*^*^_*2*_*(t)* is the apparent transverse relaxation rate, *TE* denotes the echo time, *S*_*0*_*(t)* is the magnetisation at TE = 0, and *n(t)* represents additive background noise. Thus, the relative change in signal is the sum of three terms: (i), the relative change in transverse magnetisation at zero echo time 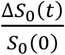; (ii), the change in relaxation rate 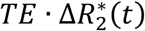; and (iii) the background noise 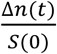 which is influenced by thermal noise, radiofrequency (RF) interference, etc. BOLD-Functional MRI experiments are typically designed to measure the relaxation term as an indirect index of functional brain activity.

There are several factors of non-neuronal origin that can influence the relative concentrations of oxygenated and deoxygenated haemoglobin in the cerebrovasculature, which can contribute to changes in the BOLD signal. These factors include heart-rate variability, respiration, and head movement (Birn, 2012; Chang et al., 2009; Friston et al., 1996; Glover et al., 2000; Parkes et al., 2018; Power et al., 2018, 2017), all of which can cause temporal correlations of the fMRI signal between brain regions. The modelling and removal of these signals continues to be an active area of research (e.g., Liu, 2017).

In addition to respiratory, cardiac and head movement factors, there are other non-neuronal factors that directly influence the BOLD signal, including the maximum oxygen carrying capacity of the blood (Gustard et al., 2003; Levin et al., 2001). This capacity is related to the amount and the fractional volume of red blood cells in the blood. Haemoglobin is the metalloprotein in red blood cells that carries oxygen from the lungs to the tissue, and returns carbon dioxide from the tissue back to the lungs. Haematocrit is the volume proportion of red blood cells to whole blood volume and is usually expressed as a percentage. Haemoglobin and haematocrit are highly correlated (haematocrit is often defined as three times the value of haemoglobin), but are not identical. Haemoglobin measures are more stable to plasma volume changes, such as in dehydration (Quinto et al., 2006). Haemoglobin and haematocrit levels vary considerably between individuals; and systematically between the sexes, with men generally showing haematocrit levels of 42-49%, and women 39-46% (e.g., Stack and Berger, 2009). Whilst haemoglobin and haematocrit levels within subjects are not thought to vary within an MRI scan session (unlike head motion or breathing rate), they are still able to influence the BOLD signal-to-noise (SNR) ratio of the acquired data. This source of variability in BOLD signal (and SNR) across subjects has the potential to influence measures of functional connectivity, particularly for analyses that examine individual variability in connectivity and differences between groups of individuals. In fact, inter-subject variation in haematocrit has been found to correlate with the degree of centrality of fMRI networks (Yang et al., 2015), to mediate the magnitude of the BOLD response in the visual cortex (Levin et al., 2001; Xu et al., 2018), and to mediate the BOLD signal intensity in other forms of oxygenation-sensitive imaging, including the cardiac BOLD response (Guensch et al., 2016).

Dependence of the BOLD signal on haematocrit level is particularly important for studies where haemoglobin and/or haematocrit may differ between study groups. Several factors have been found to correlate with haemoglobin differences, including sex (Rushton and Barth, 2010), age (Zauber and Zauber, 1987), race (Dutton, 1979), hydration level (Guensch et al., 2016), stress levels (Jern et al., 1989; Muldoon et al., 1995; Patterson et al., 1995), body temperature (Thirup, 2003), sleep apnoea (Choi et al., 2006), cardiovascular health (Jin et al., 2015), and testosterone administration (Drinka, 2013). These factors influence haemoglobin, and so may also indirectly influence fMRI functional connectivity analyses between groups and confound the results and interpretation of the findings.

In this study, we examined the potential for haemoglobin to be a confounding variable in functional connectivity analyses in a group of healthy elderly individuals. Ageing is associated with a number of physiological changes that may impact the BOLD signal (Aanerud et al., 2012), and inter-individual variability in haemoglobin levels increases across the lifespan (Hawkins et al., 1954). We specifically aimed to test whether between-group haemoglobin differences could systematically bias between-group differences in functional connectomes, and whether individual variation in haemoglobin could impact the relationships between functional connectivity and measures of cognition in group studies. We predicted that groups selected by haemoglobin levels would exhibit differences in both functional connectivity and connectivity-cognition relationships.

Given the significant known differences in haemoglobin between the sexes (Rushton and Barth, 2010), analyses were performed separately for men and women. The cohort was additionally split into two groups, divided into the lower 50% and upper 50% of individuals based upon haemoglobin values. We tested for differences in cognitive performance between the two groups to ensure functional connectivity differences were not due to differences in cognitive performance. We compared the functional connectivity matrices between the groups to identify whether there were effects of haemoglobin on the strength of global or regional functional connectivity measures. We examined whether the effect was global, and therefore an intrinsic property of the BOLD signal, or anatomically localized to specific brain networks, especially those networks with relatively high venous cerebrovascular vessel density. Finally, we examined the effect of haemoglobin variability on connectivity-cognition relationships, by correlating functional connectivity with cognitive performance and testing whether the correlations differed between the two haemoglobin groups. We hypothesised that significant differences would be observed between functional connectivity of the upper and lower haemoglobin groups, that these differences would persist in connectivity-cognition analyses, and could produce potentially spurious findings between the connectivity matrices of two otherwise comparable groups of healthy people.

## 2. Methods

This data was acquired by the ASPREE Investigator Group, under the ASPREE-Neuro sub-study. We refer the reader to the ASPREE-Neuro clinical trial protocol paper (S. Ward et al., 2017) for full study parameters. The data that support the findings of this study are available from ASPREE International Investigator Group, but restrictions apply to the availability of these data, which were used under license for the current study, and so are not publicly available. Data are however available from the authors upon reasonable request and with permission of ASPREE International Investigator Group (https://aspree.org).

All methods for the ASPREE-Neuro clinical trial were reviewed by the Monash University Human Research Ethics Committee, in accordance with the Australian National Statement on Ethical Conduct in Human Research (2007).

### 2.1 Participants

Data formed part of the baseline (i.e., premedication) cohort of the ASPREE trial, a multicentre, randomised double-blind placebo-controlled trial of daily 100mg aspirin in 19,000 healthy community dwelling older adults in Australia and the United States. Inclusion and exclusion criteria for the ASPREE trial have been published previously (Group, 2013). Participants were eligible in Australia if aged 70-years and over, no past history of occlusive vascular disease, atrial fibrillation, cognitive impairment or disability, were not taking antithrombotic therapy, and did not have anaemia or diagnosis of a conditions likely to cause death in the following five years. Baseline characteristics of the full ASPREE sample have been reported previously (McNeil et al., 2017). The current study used data from 518 participants (age=73.8±3.5, 248 females) from the ASPREE-Neuro sub-study (Ward et al., 2017). At study entry, ASPREE-Neuro participants had a Modified Mini Mental Status Examination (3MS) (Teng and Chui, 1987) score of at least 78/100. All MRIs used in this study were acquired before study medication was taken.

### 2.2 Procedure

Full protocol details are available in the ASPREE-Neuro sub-study protocol (Ward et al., 2017). Here, we include haemoglobin, cognitive and MRI data from the baseline time-point, prior to the administration of study medication.

#### 2.2.1 Haemoglobin

Fasting blood was collected at a lifestyle profile and screening assessment and sent to a pathology laboratory for testing. Haemoglobin was measured in g/dL. To comply with the ASPREE trial inclusion criteria (ASPREE Investigator Group, 2013), individuals were screened from study entry if their haemoglobin was below 11g/dL for females or 12g/dL for males.

The cohort was separated into low-haemoglobin (‘low-Hb’) and high haemoglobin (‘high-Hb’) groups using a median split (lower and upper 50%) separated by sex. This resulted in four groups: low-Hb female, low-Hb male, high-Hb female, and high-Hb male. For females, the median was 13.8, and for males the median was 14.9. Histograms of haemoglobin distribution in this sample are shown in Supplementary Figure 1.

#### 2.2.2 Cognitive tests

The five cognitive tests used were: (i) single-letter controlled oral word association test (COWAT) (Ruff et al., 1996); (ii) colour trails test (D’Elia, 1996); (iii) predicted score derived from the modified mini-mental state examination (3MS); (iv) symbol digit modalities test (SDMT) (Smith, 1982); and (v) the Victoria Stroop test (Troyer et al., 2006). Performance on each test was normalised (z-scored) separately for females and males.

#### 2.2.3 Imaging

MRI data were acquired using a 3T Skyra MRI scanner equipped with a 32-channel head and neck coil (Siemens, Erlangen, Germany). In this study, we used resting-state BOLD-fMRI and T1-weighted structural MRI data. The fMRI protocol was an eyes-open resting-state multi-band EPI sequence (multiband factor=3, TE=21ms, TR=754ms, voxel=3.0mm isotropic, matrix 64×64, slices=42). A T1-weighted MPRAGE scan was acquired for registration (TE=2.13ms, TR=2300ms, TI=900ms, voxel=1.0mm isotropic, matrix=240×256×192, flip angle=9°).

fMR images were corrected for geometry distortions (FUGUE) and brain extracted (BET) (Smith, 2002). Intra-scan movements were corrected using 3dvolreg; both high frequencies (above 0.1Hz) and the temporal mean (and first- and second-order polynomials) were removed from each voxel’s time series using 3dTproject (AFNI) (Cox, 1996). Filtered images were entered into a first-level independent component analysis with automatic estimation of the number of components using MELODIC (Beckmann et al., 2005). All the extracted ICA maps were then automatically labelled by FSL-FIX (Salimi-Khorshidi et al., 2014), which was previously manually trained on 25 random subjects. Temporal trends from noise-labelled ICA components were linearly regressed out of the 4D MR images using ordinary least squares (OLS) regression as implemented in FSL FIX. The cleaned images were then normalized to the MNI template (2mm isotropic resolution); the first volume of the EPI time series was registered to the T1-weighted image using linear registration (with 6 degrees of freedom). Each T1 was then non-linearly registered to the MNI template using the symmetric normalization algorithm in ANTs (Avants et al., 2008). The brain was parcellated into 82 regions using the Desikan-Killiany atlas (Desikan et al., 2006; Fischl et al., 2002).The last step was to apply all the transformations from the previous two points to the 4D cleaned file. Finally, the normalized cleaned file was smoothed with a 5mm FWHM Gaussian kernel.

#### 2.2.4 Calculation of functional connectivity matrix

fMRI timeseries were extracted from each of the 82 Desikan-Killiany regions for each individual. An individual functional connectivity matrix was calculated as an 82×82 correlation matrix, formed by calculating temporal correlations (Pearson) between each pair of regions. A single entry in the correlation matrix is referred to as an edge in the connectivity matrix; each region denotes a node. Group-level connectivity matrices were computed by averaging edges across subjects within groups.

#### 2.2.5 Group-level assessment of haemoglobin on the functional connectivity matrix

All analyses were performed separately for males and females to mitigate the confounding effects of haemoglobin differences associated with sex (Rushton and Barth, 2010). The effect size of functional connectivity differences between two subgroups for each sex (median split on haemoglobin values) was quantified by calculating Cohen’s d (Cohen, 2013) for each edge in the 82×82 correlation matrix. Two subgroup connectivity matrices for each sex were calculated taking the mean edge-weight across individuals in the low-Hb and high-Hb subgroups (divided by the median haemoglobin value) and the subgroup-level matrices were compared using Cohen’s d-statistic

#### 2.2.6 Origins of the haemoglobin influence

Further analysis was performed to assess whether the effect of haemoglobin on the functional connectivity matrix was due to neurovascular effects, and potentially biased towards the neuroanatomical location of the draining blood vessels. A general linear model for each edge in the matrix was fit to infer a relationship between haemoglobin (Hb) and functional connectivity (FC), covarying for age.

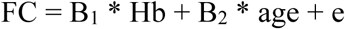

Haemoglobin values were normalised (z-scored) using a separate mean and standard-deviation for females and males to avoid previously reported biases (Rushton and Barth, 2010) prior to fitting the linear models. This model was fit using the whole cohort, i.e., not split into high and low haemoglobin groups.

The distribution of the t-scores for the linear coefficients (B_1_) and explained variance of the entire model (R-squared) were compared to a null model where the Hb, FC and age data were randomly permuted, and the models refit at each edge. The distribution for the null model was the composite of t-scores and explained variance from 500 permutations. We calculated an edge-wise p-value from the number of permutations with higher t-score values.

To determine whether the proximity of brain regions to cerebrovenous vasculature influenced the relationship between haemoglobin and functional connectivity, the location of the strongest associations were compared to an atlas of the cerebral veins (Ward et al., 2018). For each sex, a ‘haemoglobin connectome’ was constructed with edges to represent the top 10% of linear coefficients (t-values). The number of edges connecting each node was then calculated (network degree) to determine which regions of the brain gave rise to the strongest haemoglobin-functional connectivity associations. These highly connected regions were then spatially compared to the probabilistic map of the location of the cerebral veins (Ward et al., 2018).

#### 2.2.7 Impact of haemoglobin on connectivity-cognition analyses

To investigate how group-differences in haemoglobin may influence brain connectivity-cognition relationships, the relationships between cognitive performance and functional connectivity were compared between the two subgroups (low-Hb and high-Hb) for each sex. Thus, in the comparison of low-Hb versus high-Hb groups within each sex, the mean haemoglobin values are the only between-group difference. The two haemoglobin subgroups for each sex had no differences in cognitive performance for either the male or female groups (see Results).

A linear model was calculated between functional connectivity (FC) and task performance (Cog), covarying for age, for each of the five tasks, for each edge in the connectivity matrix.

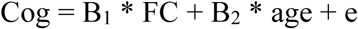

The linear coefficients (B_1_) of the lower-Hb and upper-Hb subgroups were compared to measure the effect of subgrouping on haemoglobin using a Cohen’s d statistic.. To assess the effect of haemoglobin, the results were compared with linear coefficients estimated from the entire cohort (low-Hb and high-Hb combined).

Functional connectivity and cognitive performance were normalised (z-scored) prior to fitting the models. All normalisation and statistical analyses were performed separately for females and males.

## 3. Results

### 3.1 Group-level assessment of haemoglobin on the functional connectivity matrix

Group-level functional connectivity matrices differed significantly between the low-Hb and high-Hb sub-groups for both men and women (t-test p<10^−10^). The size of this effect at a global level was small (Cohen’s d men=0.17 women=0.03). In males, the edge-weights of the functional connectivity matrix in the high-Hb subgroup were consistently higher than in low-Hb subgroup (Figure 1B) with effect sizes as high as 0.4. For men, 88% of the edge-weights were higher in high-Hb compared to low-Hb subjects, with 12% having higher edge-weights in the low-Hb compared to the high-Hb subjects. In females, there was less systematic bias between the subgroup matrix edge weights with 59% higher in the high-Hb compared to the low-Hb subgroup, with 41% higher in the low-Hb subgroup than high-Hb subgroup. The strength of the global haemoglobin effect between sex subgroups was less in females compared to males (Figure 1A).

**Figure 1.**
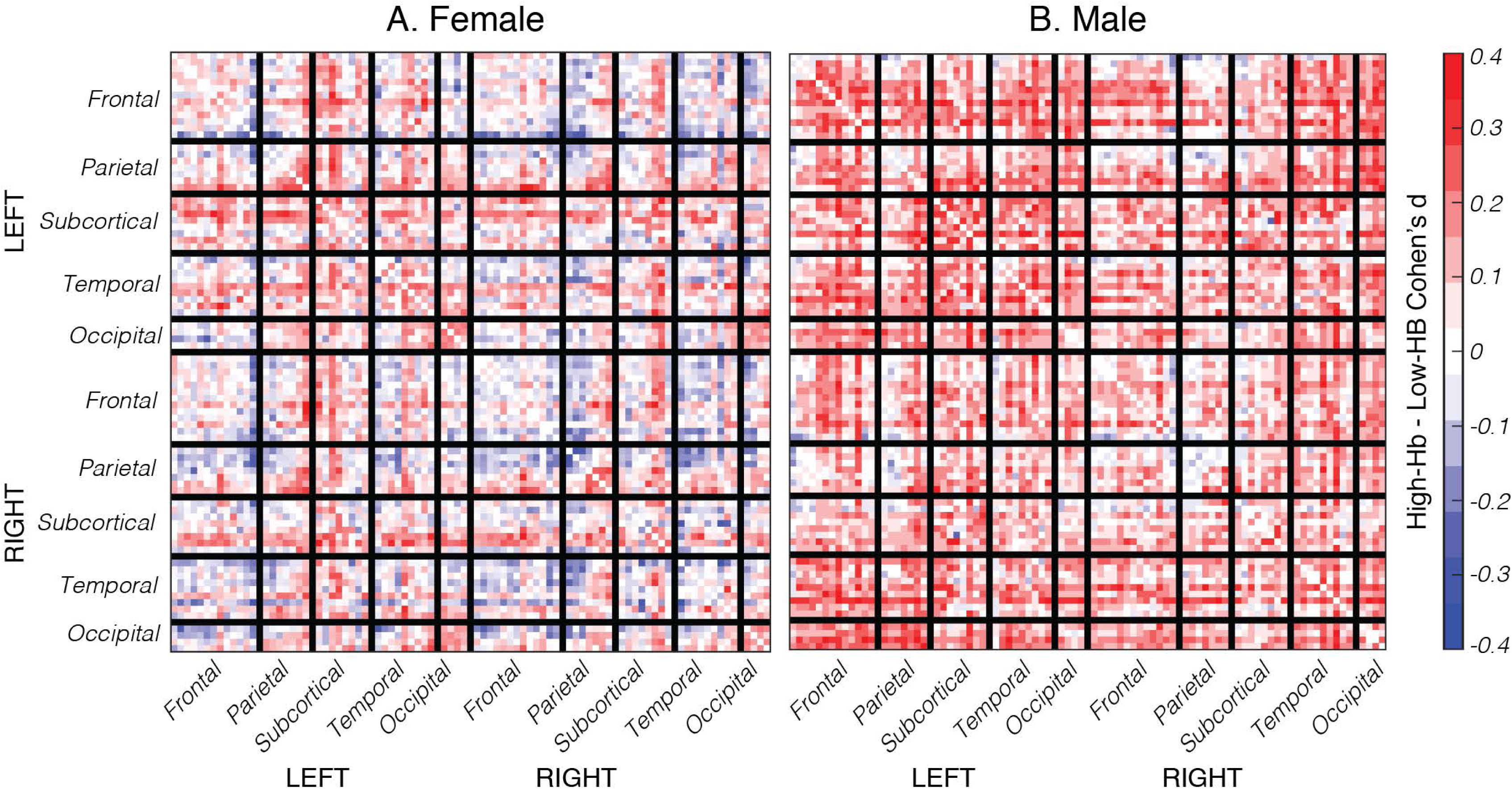
The effect of haemoglobin on resting-state functional connectivity for (A) females and (B) males. Values are Cohen’s d effect sizes of the difference between high-haemoglobin and low-haemoglobin groups. Abbreviation: Hb, haemoglobin

### 3.2 Origins of the haemoglobin influence

Linear coefficients between haemoglobin and edge-weights in the functional connectivity matrix were calculated and compared to a null distribution of randomly permuted haemoglobin, functional connectivity and age data. In men, the linear coefficients (Figure 2B.i.) and explained variance (Figure 2B.ii.) were linearly biased relative to the randomly permuted dataset. The likelihood of this bias is depicted in Figure 2B.ii and compared to the expectation of a null model. These results demonstrate that the results are highly unlikely to be false-positives.. The distribution of linear coefficients was biased positively in the haemoglobin model, relative to the null data. This was not the case in the female group, where the distribution of the linear coefficients between haemoglobin and functional connectivity edge-weights overlapped the null distribution (Figure 2A).

**Figure 2:**
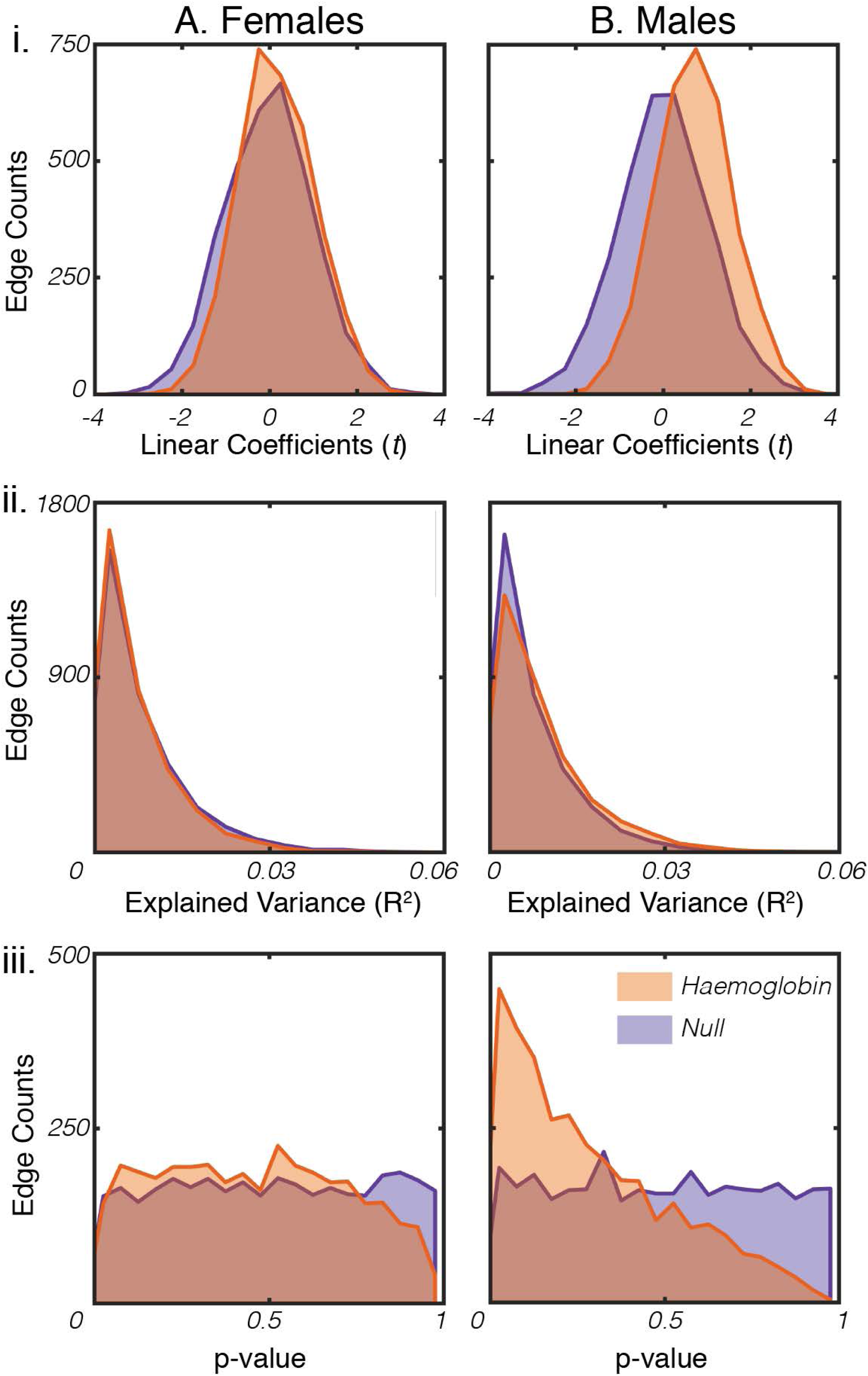
Correlation between resting-state functional connectivity and haemoglobin values in (A) Females and (B) Males. Upper plots (i.) show distribution of linear coefficients (t-values). Middle plots (ii.) show distribution of explained variance, and lower panel (iii.) shows distribution of p-values. Each panel compares the distribution of obtained values to a null distribution, calculated by randomly permuting haemoglobin values between subjects and refitting the identical model.

To determine if the haemoglobin influence on functional connectivity values was related to the proximity to cerebral veins, a map of the highest haemoglobin functional connectivity associations was compared with an atlas of the cerebral veins (Ward et al., 2018). Figure 3 shows the spatial map of the top 10% *t* coefficients, and the probabilistic location of the major draining veins (Ward et al., 2018). The strongest correlations between haemoglobin and functional connectivity were not found in close proximity to cerebral veins. Note that spatial maps at different thresholds are shown in Supplementary Figure 2 and are consistent with these results, indicating that this result is robust to different thresholds. Visual inspection of Figure 3 suggests there was a trend towards higher associations in the right hemisphere in both males and females, most pronounced in the sub-cortical regions.

**Figure 3:**
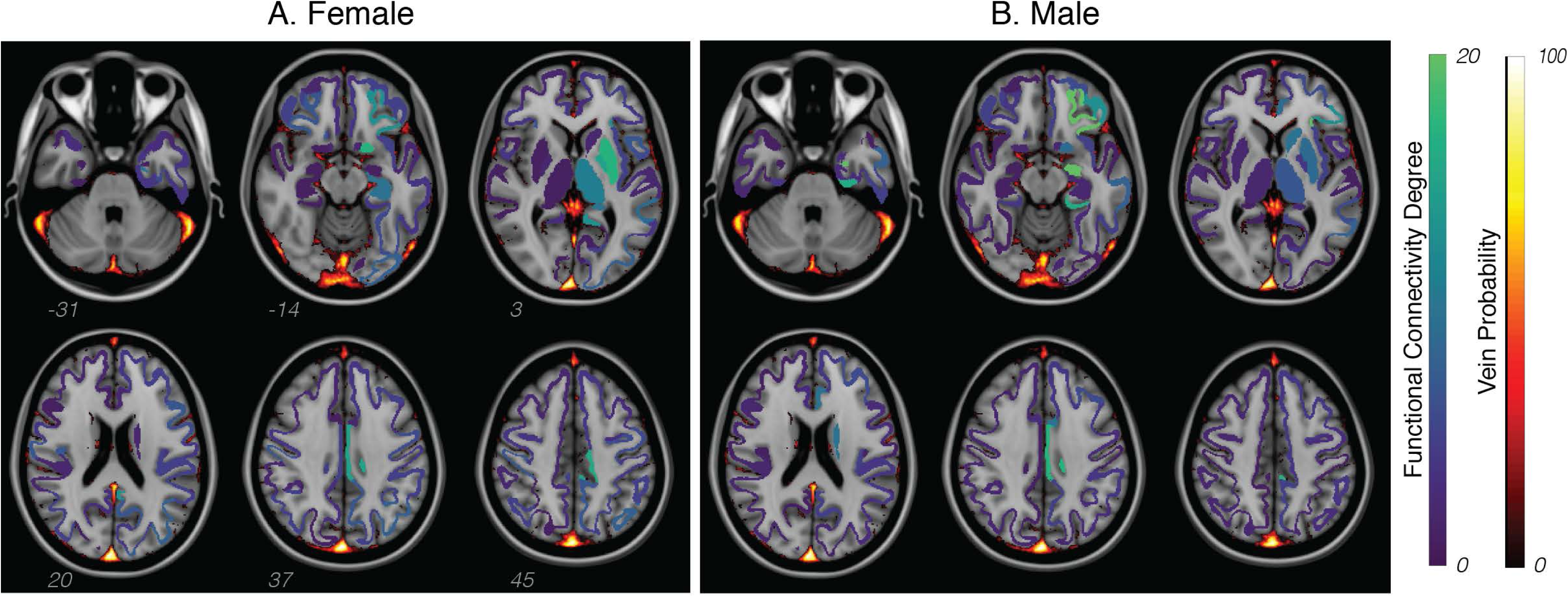
Spatial representation of regions strongly influenced by haemoglobin in resting-state functional connectivity (region degree, blue-green colour bar), and the probabilistic location of the major draining veins (see Ward et al., 2018; red-yellow colour bar), for (A) females and (B) males. Degree is defined by the number of edges connected to a region with correlation in the 90^th^ percentile. Supplementary Figure 2 show results at different thresholds and standardised to a t-distribution.

### 3.3 Impact of haemoglobin on connectivity-cognition analyses

There were no significant differences in cognitive performance between the low-Hb and high-Hb subgroups (t-test p>0.36 for all tests and both sexes). Supplementary Figure 3 shows box-plots for low-Hb and high-Hb groups for each sex.

Linear coefficients of cognitive test performance and edge-weights in the functional connectivity matrix were compared between the low-Hb and high-Hb subgroups. In both females and males, the two sets of coefficients were significantly different in four of the five cognitive tests (Table 1).

**Table 1:**
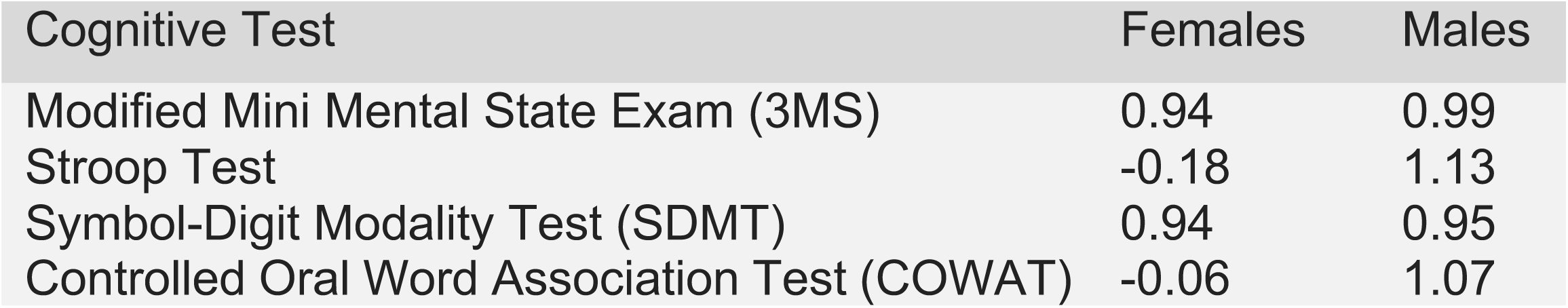
The size of the effect (Cohen’s d) between haemoglobin sub-groups on cognition-connectivity relationships.

Linear coefficients between functional connectivity and cognitive performance were consistently lower in the low-Hb subgroup compared to the high-Hb subgroup (Figure 4). The effect was present in both males and females. Matrix edges that showed no significant correlation between cognitive performance and functional connectivity (where the black line approached the zero line) still demonstrated a haemoglobin-related bias.

**Figure 4:**
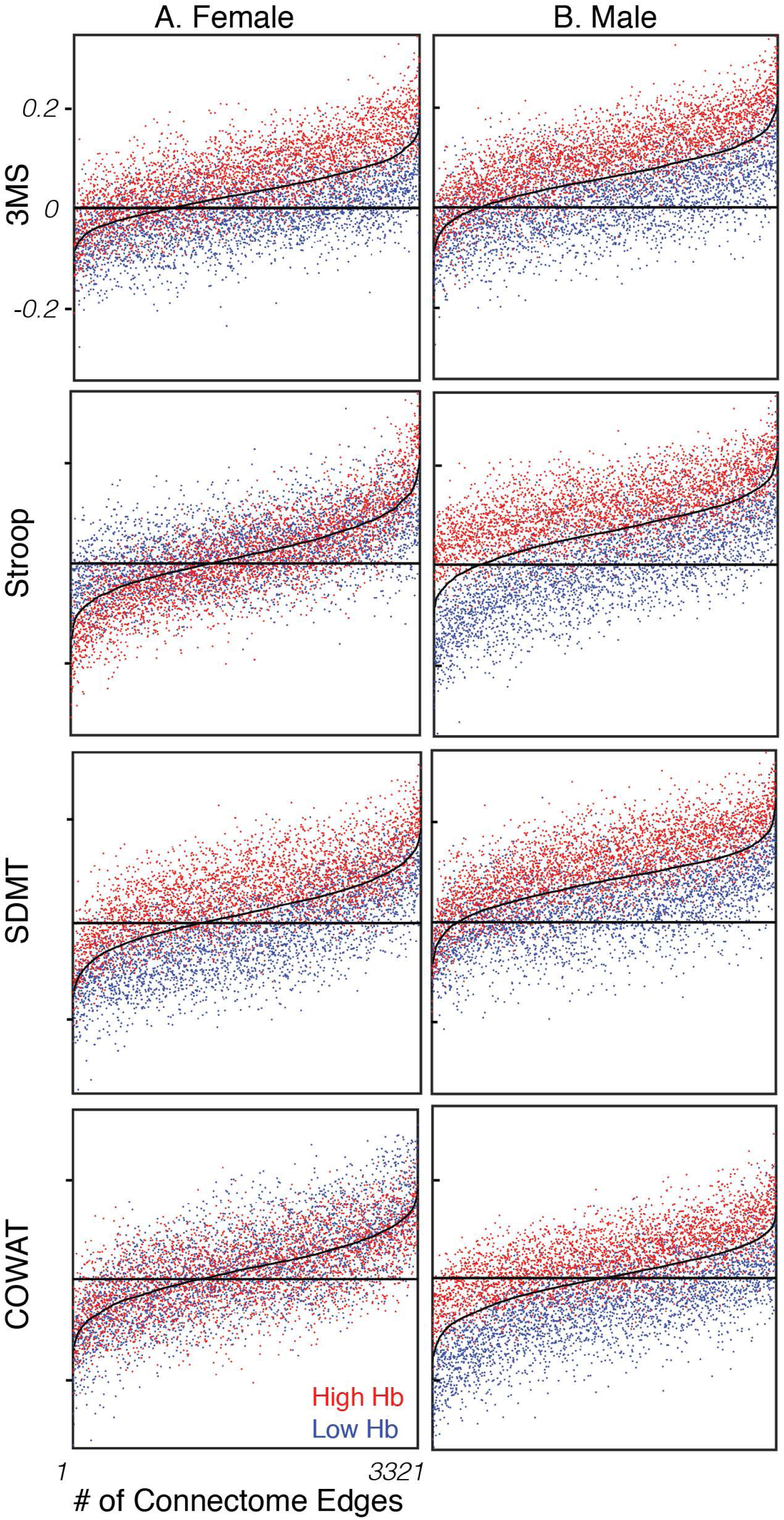
Correlation between resting-state functional connectivity and cognitive test performance in five domains, (top-bottom) 3MS, Stroop, SDMT, COWAT. Each dot denotes an edge in the functional connectivity matrix correlated with the cognitive test score. The black line shows the relationship between the cognitive score and functional connectivity calculated for the entire cohort. The blue dots show the relationship between the cognitive score and functional connectivity for the low haemoglobin group. The red dots show the relationship between the cognitive score and functional connectivity for the high haemoglobin group. Abbreviations: 3MS, modified mini-mental state exam; SDMT, symbol-digit modalities test; COWAT, controlled word association test; Hb, haemoglobin.

## 4. Discussion

The BOLD signal originates in the blood, so it is therefore not surprising that group and individual differences in haemoglobin levels are an important confounding variable in BOLD-fMRI-based studies of functional connectivity. The results of this study demonstrate that the confounding effect of variability in haemoglobin values is widespread across brain regions, differs substantially between the sexes, and strongly influences functional connectivity-cognition relationships. The effect of haemoglobin on functional connectivity measures was widespread across brain regions in males without particular neuroanatomical specificity. In females, the effect was weaker than that in the males: it varied across the brain, with subcortical regions in particular showing higher functional connectivity in the high-Hb subgroup compared to the low-Hb subgroup. Females also showed regionally specific higher functional connectivity in the low-Hb compared to the high-Hb subgroup, particularly in parietal and temporal regions. These results demonstrate that BOLD-fMRI functional connectivity analyses are confounded by haemoglobin differences, especially in studies aiming to compare groups of individuals, or between sexes.

In addition to the significant effect of haemoglobin on functional connectivity, we also found that cognition-connectivity relationships were substantially impacted by haemoglobin levels. These results showed that there are systematically higher correlations of cognitive measures with resting-state connectivity for individuals with higher haemoglobin levels, and that this effect is spatially-non-specific, and occurs across the brain. Thus, the influence of haemoglobin variability is not confined to any one individual cognitive measure, or any single brain region (e.g., close to cerebral veins). These results suggest that care should be taken when interpreting connectivity-cognition relationships calculated at the group level, that do not account for individual variability in haemoglobin levels.

The BOLD-fMRI signal relies upon the magnetic properties of haemoglobin, which itself is closely related to haematocrit: the proportion of red blood cells in the blood. When neuronal activity within a brain region increases, the additional neuronal metabolic activity results in an increased local energy requirement, which is reflected as an increase in the regional cerebral metabolic rate of oxygen consumption (CMRO_2_) (Buxton and Frank, 1997; Glover, 2011). The consumption of oxygen results in an early elevation of deoxygenated haemoglobin in the region. Following neuronal activity, the increased metabolic requirement results in an increase in cerebral blood flow (CBF) to the region to restore the local O_2_ levels. However, the CBF to the region increases by a larger proportion than required to satisfy the increased CMRO_2_ requirement. Consequently, the local concentration of deoxyhaemoglobin decreases, leading to an increase in the MR T2* signal. In sum, combined changes in CBF, cerebral blood volume (CBV) and CMRO_2_ are reflected in changes in local deoxygenated haemoglobin. The fMRI acquisition sequence is tuned to detect changes in the apparent transverse relaxation rate (T2*), which is sensitive to the amount of deoxygenated haemoglobin in the blood (Buxton et al., 2004; Liu, 2017). Therefore, individual differences in the proportion of haemoglobin and red blood cells are evident as individual differences in BOLD SNR. Our results complement the previous studies showing that BOLD signal intensity is mediated by individual haemoglobin levels (Levin et al., 2001; Guensch et al., 2016; Xu et al., 2018).

In the majority of BOLD-fMRI experiments, the measurement of interest is the relative change in the BOLD signal across time (during rest, in response to a task, etc.). In other words, the change in relaxation rate 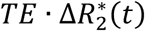 is interpreted as an indirect index of functional brain activity (Liu, 2017). As the relaxation rate in BOLD-fMRI depends on the total amount of deoxygenated haemoglobin in the blood, individual differences in haemoglobin/haematocrit will specifically influence the relaxation term 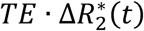. In other words, the very signal of interest in BOLD-fMRI experiments is influenced by an individual’s haemoglobin level. Notably, an individual’s haematocrit level also influences the viscosity of the blood, with higher haematocrit levels significantly slowing the rate of blood flow throughout the body (Stack and Berger, 2009). Differences in CBF influences the number of spins that flow into a voxel (Gao and Liu, 2012), which affects the magnetisation term 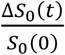 of the BOLD signal. Individual differences in haemoglobin may therefore also indirectly influence the magnetisation term of the BOLD signal. Previous studies that have aimed at minimising the effect of noise and confounds on the BOLD signal have focused on factors that vary across time, including cardiac noise, respiratory noise, motion, and low frequency drifts and fluctuations (Liu, 2016). Individual differences in haemoglobin/haematocrit have not been considered as an important confound, possibly because this value does not vary for an individual during a typical scan duration. This appears to be especially so for studies of functional connectivity, which by definition examine relationships between signals that vary across time.

The relationship between BOLD-fMRI and individual differences in haematocrit has been investigated in a number of small cohort studies. Yang et al. (2015) found only a modest relationship between haematocrit variability and summary measures of functional connectivity in a sample of 45 healthy younger (mean age 29yrs; s.d. 10.8) adults. Degree of centrality, fractional ALFF (amplitude of low frequency fluctuations) and voxel-mirrored homotopic connectivity showed discrete clusters of significant association with haematocrit, in non-overlapping regions. Resting-state networks estimated with dual regression showed more widespread relationship with between-subject variation in haematocrit. They concluded that the impact of haematocrit variation in BOLD signal is modest and regionally-specific. In another sample of 13 healthy middle-aged (mean age 35yrs; s.d. 7) adults, Xu et al. (2018) found that 22% (i.e., R^2^=0.22) of the inter-subject variability of the BOLD response in the visual cortex to flickering checkerboard stimulus was associated with haematocrit, consistent with previous studies of haematocrit and the BOLD signal (R^2^=0.23; Gustard et al., 2003; R^2^=0.28, Levin et al., 2001). When the BOLD response was normalised by the haematocrit, the BOLD signal coefficient of variation reduced by 16% leading Xu et al. (2018) to conclude that normalisation of the BOLD signal by individual haematocrit levels is an important step for enhancing the detection power of BOLD-fMRI studies.

The current sample is substantially larger than the previous studies of the influence of haemoglobin on BOLD-fMRI measures (Yang et al., 2015; Xu et al., 2018), with a sample size of 518 subjects. Like Xu et al. (2018) our results suggest that individual differences in haemoglobin have a substantial influence on BOLD-fMRI measures of functional connectivity and functional connectivity-cognition relationships. Notably, Xu et al. (2018) used a task-based (stimulus-evoked) paradigm, suggesting that the influence of haemoglobin on fMRI measures are not limited to resting-state paradigms alone. In contrast to Yang et al. (2015), we found that haemoglobin effects were widespread, spatially non-specific in men; and widespread and spatially-variable in women. Unlike the previous studies, our sample were older adults within a very narrow age range (73.8±3.5yrs), with no major health problems and who passed stringent criteria for inclusion in a clinical trial (Ward et al., 2017).

Ageing is associated with a number of physiological changes that may impact the BOLD signal, including changes in oxygen extraction fraction, CMRO_2_, and CBF (Aanerud et al., 2012). Haemoglobin declines with age in men, and increases with age in women (Bäckman et al., 2016; Cruickshank, 1970), such that the values for men and women become more similar in older age, but still significantly different (note however this finding is not ubiquitous, with findings of declines in haemoglobin levels for both sexes (Salive et al., 1992)). Haematocrit does not seem to change post-menopause compared to pre-menopause (Amin et al., 2004; Koons et al., 2019), however CMRO_2_ has been found to change which may pose additional challenges for fMRI studies in older females (Peng et al., 2014). Although we do not have data on the menopausal status of the women in this study, it is notable that all participants were aged over 70-years, and the mean age of menopause in Australia is 51.3-years (Davis et al., 2015). As such, the majority of the women in this sample are likely to be post-menopausal. An important direction for future research is to examine how menopause influences BOLD physiology.

Haemoglobin variations are known to coincide with cerebral blood flow changes (van de Veen et al., 2014). It is possible that the correlations observed in this work are not a direct consequence of haemoglobin but a composite effect of blood flow, blood volume, and vascular reactivity, all of which may be related to haemoglobin levels. Furthermore, these effects may both directly affect the BOLD signal and modulate the haemodynamic response, providing multiple mechanisms for further exploration.

Taken together, our results provide evidence that individual differences in haemoglobin/haematocrit are an important confounding variable in functional connectivity and functional connectivity-cognition analyses. The results suggest that it is possible that previous studies of sex differences in functional connectivity and connectivity-cognition relationships that did not control for haemoglobin may be confounded, at least in ageing (e.g., Jamadar et al., 2018). Our results are consistent with some (Xu et al.; Guensch et al., 2016), but not other (Yang et al.) results in healthy younger adults. Given the differences in haemoglobin concentration over the lifespan (Backman et al. 2016, Cruikshank, 1970; Salive et al., 1992), future studies should explore if the effect of haemoglobin on the BOLD-fMRI signal varies with age.

A number of approaches exist for obtaining measures of haemoglobin for inclusion in BOLD-fMRI analyses. One approach, as used in this study, involved drawing venous blood and laboratory measurement of full blood parameters. However, smaller devices that provide point-of-care metrics of haemoglobin and haematocrit are now available (Nkrumah et al., 2011; (Avcioglu et al., 2018; Singh et al., 2015). These devices require finger-prick samples of capillary blood, are quick, easy to obtain, and relatively non-invasive. Comparisons of point-of-care devices for measuring haemoglobin have been performed (Avcioglu et al., 2018; Singh et al., 2015). Avcioglu et al. showed that the point-of-care measurement of haemoglobin was correlated with venous haemoglobin measures analysed with gold-standard cell analysers, with R^2^=0.825. Alternative approaches include image-derived estimates of haematocrit using the dependence of the T1 relaxation rate in venous blood (Li et al., 2017), which takes around two minutes to acquire, and requires additional positioning of the subject to image the chest for complete inversion of the inflowing arterial blood (Li et al., 2017). Image-derived estimates of haematocrit were found to correlate strongly (R^2^=0.91) with laboratory based measures of complete blood count (Xu et al., 2018). In sum, the ease and cost of devices for acquiring haemoglobin or haematocrit metrics should not be a barrier for most groups acquiring BOLD-fMRI data.

One limitation of the current study is the known regional variation of haematocrit throughout the brain. Calamante et al. showed that local measures of capillary haematocrit can be obtained by combining multiple MRI sequences, namely arterial spin labelling and dynamic susceptibility contrast (Calamante et al., 2015). This approach could be used in future studies to compare the influence of haematocrit variation on regional BOLD fMRI signal characteristics, to potentially provide a neuroanatomical map of the degree of haematocrit influence on the estimation of functional connectivity metrics. Given the known variation in haematocrit values across the body (Mchedlishvili & Varashvili, 1987), brain image-derived metrics of haematocrit may be ultimately be the best method to control for haemoglobin effects in BOLD-fMRI analyses. It is also important to note that resting-state functional connectivity is sensitive to differences in pre-processing pipelines (e.g. Parkes et al., 2018) and resting-state condition (e.g., eyes open, eyes closed, fixation; Patriat et al., 2013). Future studies should confirm the influence of haemoglobin on estimates of functional connectivity under different conditions and analysis pipelines. Finally, the demographic characteristics of the current sample should be noted when interpreting these results. To comply with the inclusion criteria for the ASPREE clinical trial (during which this data was acquired), the lower limit of haemoglobin values was restricted to above 11g/dL for females and 12g/dL for males. This restricted range may have led to us underestimating the effect of haemoglobin variability on functional connectivity (Goodwin & Leech, 2006). The ASPREE clinical trial is also predominantly white (91%; McNeil et al., 2017), and mean haemoglobin values are known to differ by race (e.g., Dong et al., 2008). As such, future work in this area should consider samples with less restricted and more representative range of haemoglobin values.

In conclusion, this study in a large sample of healthy older adults demonstrated that individual variability in haemoglobin has a substantial influence on functional connectivity and functional connectivity-cognition relationships. The effect of haemoglobin was widespread, differed substantially between the sexes, and strongly influenced functional connectivity-cognition relationships. In males the effect of haemoglobin on functional connectivity measures was widespread across brain regions whereas in females the effect varied across the brain including in subcortical regions. Acquisition of haemoglobin/haematocrit measures are readily available and future BOLD-fMRI functional connectivity and connectome studies should control for haemoglobin as a confounding variable, especially in studies aiming to compare groups of individuals, compare sexes, or examine connectivity-cognition relationships.

## 5. Acknowledgements

PGDW & SDJ conceived the research question. PGDW, SO, AA, SF, ERO conducted the analysis. SDJ & PGDW wrote the first draft of the manuscript. SDJ & PGDW wrote the revision, with GFE edits. All authors contributed to manuscript preparation and review.

PGDW, SDJ, ERO, AF & GE are supported by the Australian Research Council (ARC) Centre of Excellence for Integrative Brain Function (CE140100007). SDJ is supported by an ARC Discovery Early Career Researcher Award (DE150100406) and NHMRC Fellowship (APP1174164).

We thank the ASPREE group for data access. ASPREE was supported by the National Institutes of Health (grant number U01AG029824); the National Health and Medical Research Council of Australia (grant numbers 334047, 1127060, 1086188); Monash University (Australia) and the Victorian Cancer Agency (Australia). The Principal ASPREE study is registered with the International Standardized Randomized Controlled Trials Register, ASPirin in Reducing Events in the Elderly, Number: ISRCTN83772183 and clinicaltrials.gov number NCT01038583. ASPREE-Neuro trial is registered with Australian New Zealand Clinical Trials Registry ACTRN12613001313729.

The authors have no competing or conflicting interests.

## Supplementary Material

**Supplementary Figure 1:**
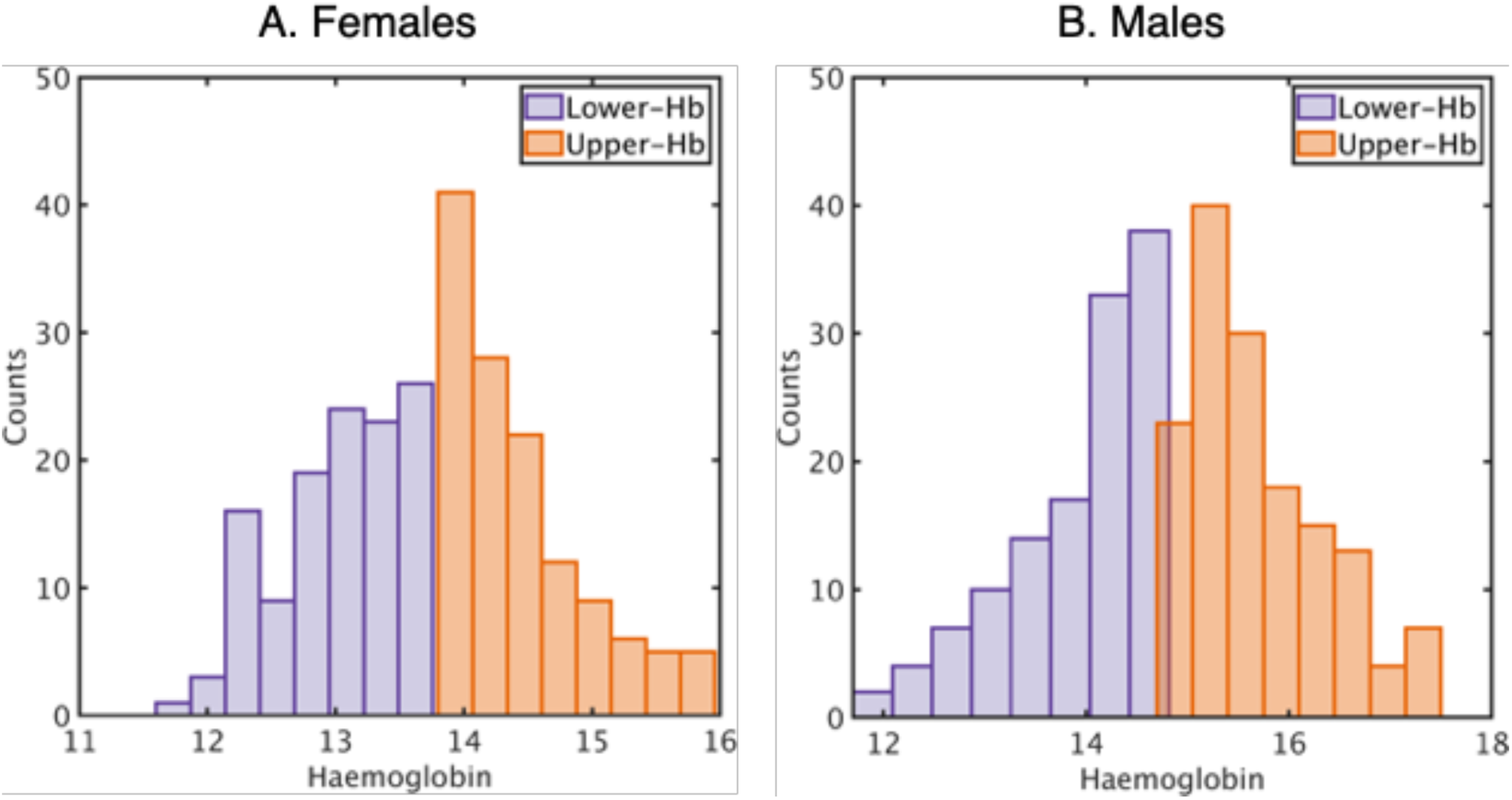
Histograms of haemoglobin distributions in this sample, for (A) females and (B) males

**Supplementary Figure 2:**
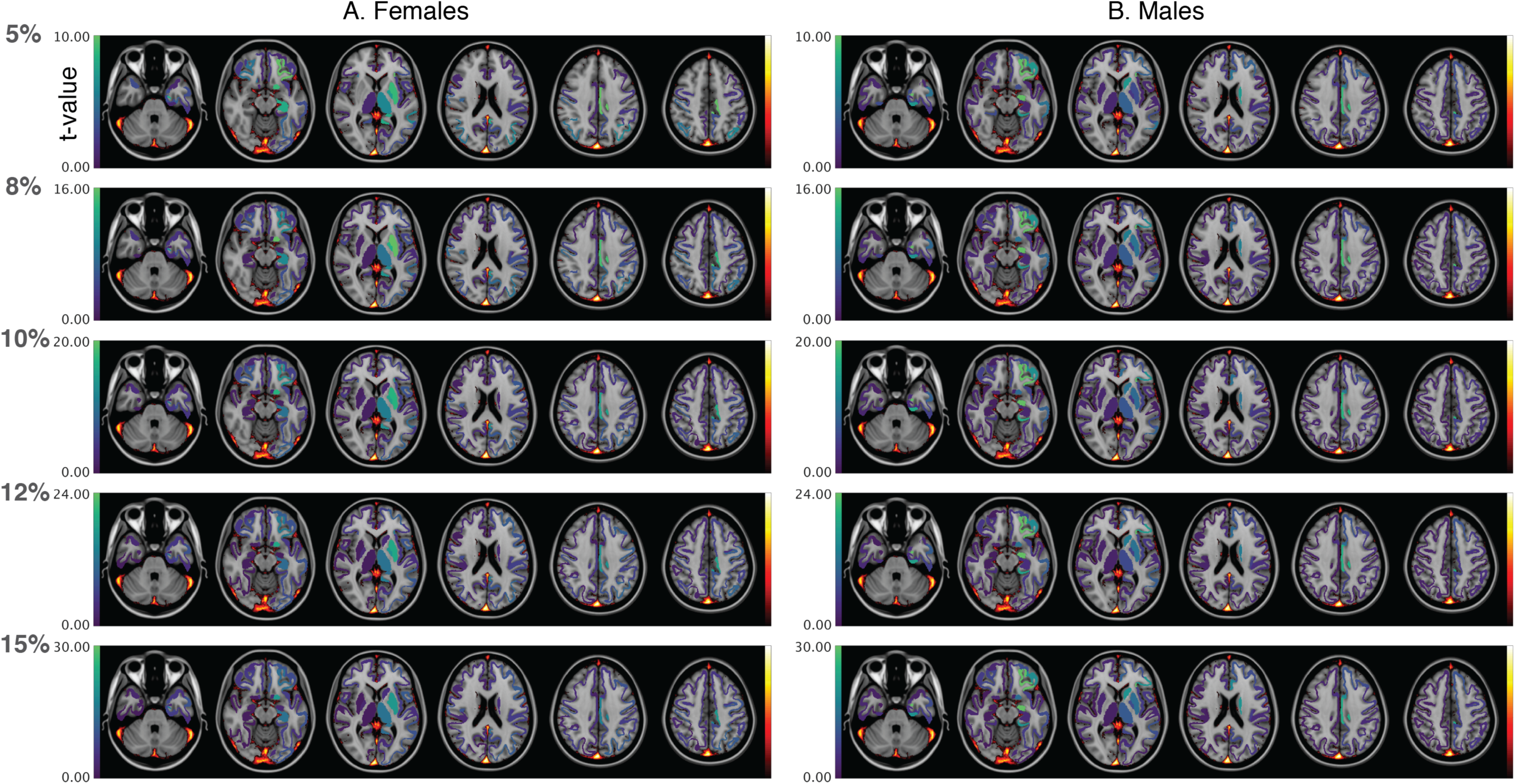
Spatial representation of regions influenced by haemoglobin in resting-state functional connectivity (region degree, blue-green colour bar) and the probabilistic location of the major draining veins. Here, haemoglobin-rsfMRI values are standardised to a t-distribution, and shown at five thresholds: top 5%, 8%, 10%, 12% and 15%.

**Supplementary Figure 3:**
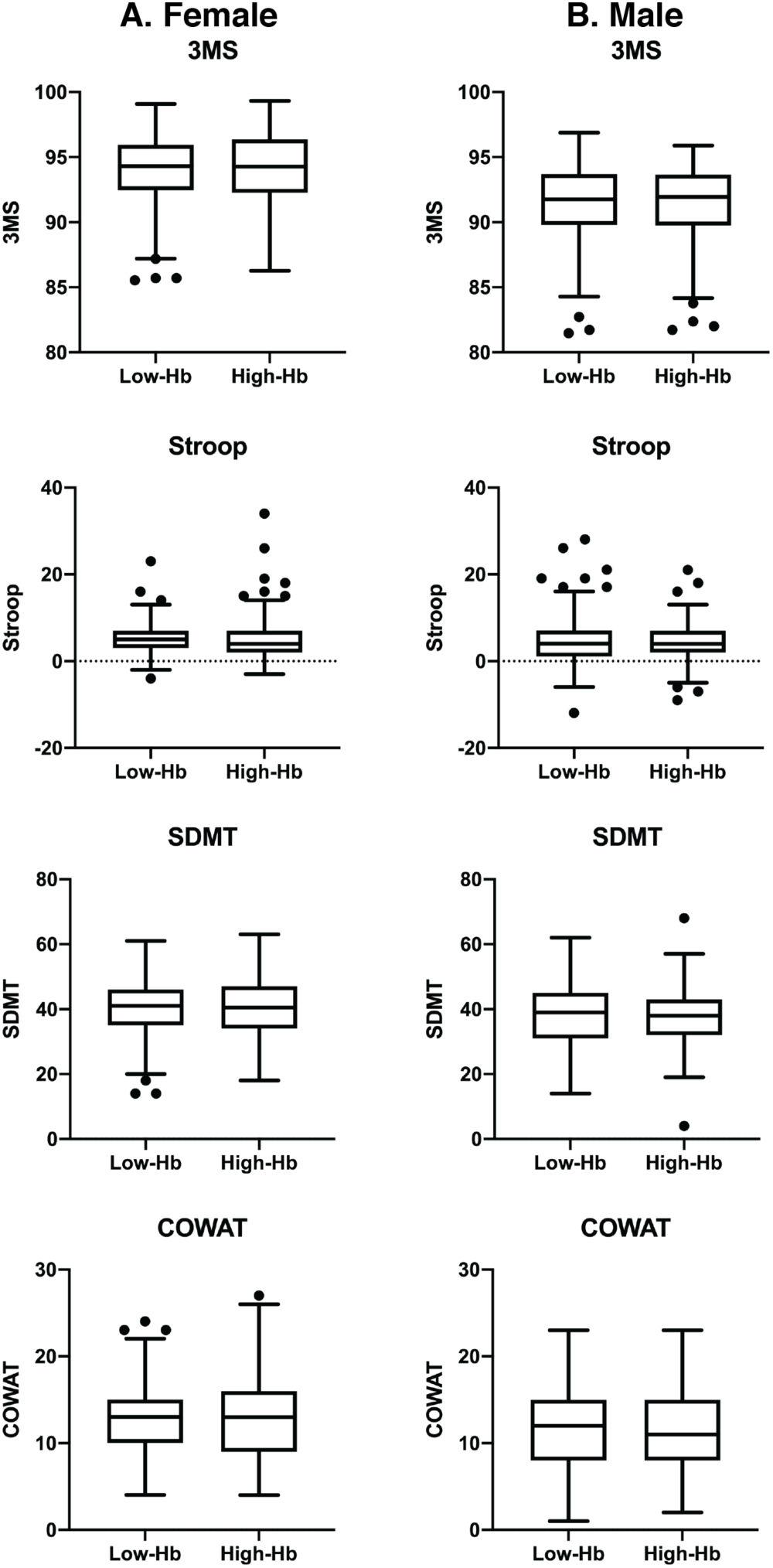
Tukey box-plots of cognitive scores for (A) females and (B) males for low haemoglobin (Low-Hb) and high haemoglobin (High-Hb) groups. Abbreviations: 3MS, Modified Mini Mental State Exam; SDMT, symbol-digit modality test; COWAT, controlled oral word association test.

